# Long noncoding RNA-mediated epigenetic regulation of auxin-related genes controlling shade avoidance syndrome in *Arabidopsis thaliana*

**DOI:** 10.1101/2023.03.06.531280

**Authors:** María Florencia Mammarella, Leandro Lucero, Nosheen Hussain, Aitor Muñoz-Lopez, Ying Huang, Lucia Ferrero, Guadalupe L. Fernandez-Milmanda, Pablo Manavella, Moussa Benhamed, Martin Crespi, Carlos L. Ballare, José Gutiérrez Marcos, Pilar Cubas, Federico Ariel

## Abstract

The long noncoding RNA (lncRNA) *AUXIN-REGULATED PROMOTER LOOP* (*APOLO*) recognizes a subset of target loci across the *Arabidopsis thaliana* genome by forming RNA-DNA hybrids (R-loop) and modulating local three-dimensional chromatin conformation. Here we show that *APOLO* is involved in regulating the shade avoidance syndrome (SAS) by dynamically modulating the expression of key factors. In response to far-red (FR) light, the expression of *APOLO* anticorrelates with its target *BRANCHED1* (*BRC1*), a master regulator of shoot branching in *Arabidopsis thaliana*. *APOLO* deregulation results in *BRC1* transcriptional repression and an increase in the number of branches. *APOLO* transcriptional accumulation fine-tunes the formation of a repressive chromatin loop encompassing the *BRC1* promoter, which normally occurs only in leaves as well as in a late response to FR treatment in axillary buds. In addition, our data reveal that *APOLO* participates in leaf hyponasty, in agreement with its previously reported role in the control of auxin homeostasis through direct modulation of *YUCCA2* (auxin synthesis), *PID* and *WAG2* (auxin efflux). We found that direct application of *APOLO* RNA to leaves results in a rapid increase in auxin accumulation that is associated with changes in the response of the plants to FR light. Collectively, our data support the view that lncRNAs coordinate the shade avoidance syndrome in *Arabidopsis thaliana* and shed light on the potential of lncRNAs as bioactive exogenous molecules. Deploying exogenous RNAs that modulate plant-environment interactions are important new tools for sustainable agriculture.

## INTRODUCTION

Plants can adapt their architecture and physiology in response to environmental conditions and endogenous signals throughout their entire life cycle. This plasticity increases the efficiency of resource capture and utilization and is a central determinant of plant fitness in variable environments (Sultan, 2010). One of the best studied cases of phenotypic plasticity in plants is the shade avoidance syndrome (SAS) (Smith, 1995). SAS is characterized by morphological and physiological responses that allow plants to avoid being shaded by neighbors and, therefore, increase the interception of sunlight in plant canopies. These responses are triggered by light cues that indicate the proximity of other plants, particularly the reduction in the ratio of red (R) to far-red (FR) radiation (R/FR ratio) that is caused by the absorption of R photons by chlorophyll-containing tissues (Casal, 2012; Ballaré and Pierik, 2017; Fiorucci and Fankhauser, 2017). Changes in the R/FR ratio are perceived by the photoreceptor phytochrome B (phyB), which controls numerous aspects of plant growth and development (Legris et al. 2019). The morphological responses that allow plants to avoid shade are manifold and depend to a large extent on the architecture and developmental stage of the species. Among the best characterized morphological responses to a reduction in the R/FR ratio are the increase in the rate of stem elongation, increased apical dominance (i.e., reduced branching) and repositioning of the leaves to adopt a more vertical orientation (leaf hyponasty) (Fernández-Milmanda and Ballaré, 2021). The combination of these responses results in a configuration of the plant shoot that increases the likelihood of intercepting photons in crowded plant stands.

Shoot branching, which relies on the capacity of axillary buds to grow and form a new branch, is frequently inhibited as part of the SAS, in order to prioritize the growth of the main stem. In *Arabidopsis thaliana*, the axillary bud outgrowth is repressed by *BRANCHED1* (*BRC1*), a gene encoding a transcription factor from the TCP class II family that is expressed in axillary buds. *BRC1* is considered a master regulator of branching because it integrates numerous endogenous and exogenous signals, including hormone balance and R/FR light ratio (Aguilar-Martínez et al., 2007; González-Grandío et al., 2013; Rameau et al., 2015).

Leaf hyponasty is a typical shade avoidance response, particularly in rosette plants such as *Arabidopsis thaliana*, and can be very important to optimize light capture in dense canopies (Pantazopoulou et al., 2017). The response is triggered by low R/FR ratios sensed by phyB, and this signal is perceived more effectively at the leaf tip (Michaud et al., 2017; Pantazopoulou et al., 2017). Leaves respond to the low R/FR signal with asymmetrical growth between the abaxial and adaxial sides of the petiole, which results in upward movement of leaf blades. The plant hormone auxin plays a central role in this response. Recent studies have shown that low R/FR ratios perceived at the leaf tip promote the accumulation of leaf tip-derived auxin in the abaxial petiole. This local auxin accumulation is responsible for the asymmetric growth that causes the hyponastic response to low R/FR, and is determined by the auxin transport protein PIN3 (Küpers et al., 2023).

At the molecular level, plant developmental plasticity during SAS depends on a wide range of gene expression regulatory mechanisms. Among them, chromatin structure dynamics determine the spatial context of gene location, accessibility and transcription (Patitaki et al., 2022). Such chromatin 3D organization depends on the action of histone modifiers, long noncoding RNAs (lncRNAs), transcription factors, and chromatin remodelling and mediator proteins. In general terms, the acetylation and phosphorylation of histone tails are associated with an open chromatin conformation. Histone methylation, such as H3K27me3 or H3K9me2, are linked to chromatin compaction and frequently characterize the epigenetic profile of silenced genes and transposons, respectively (Rodriguez-Granados et al., 2016). Interestingly, lncRNAs participate in histone modification dynamics and chromatin compaction by recruiting or decoying specific proteins to/from target regions. Moreover, it has been shown that lncRNAs can directly or indirectly affect gene activity by fine-tuning chromatin 3D loop formation (Fonouni-Farde et al., 2022).

In the model species *Arabidopsis thaliana*, the lncRNA *APOLO* recognizes in *cis* its neighbor gene *PID* and a subset of target genes in *trans* through sequence complementarity and R-loop (DNA-RNA duplex) formation. As a result, *APOLO* modulates local chromatin 3D conformation and gene transcriptional activity (Ariel et al., 2014; Ariel et al., 2020). Some of these are auxin-responsive genes and *APOLO* is induced by auxin (Ariel et al., 2014). As SAS responses are often controlled by auxin, we wondered about the role of *APOLO* in SAS.

In this study we show that *APOLO* is dynamically modulated by a low R/FR ratio and participates in SAS. In axillary buds, *APOLO* directly regulates the transcription of the master regulator *BRC1* by controlling a local chromatin loop encompassing the *BRC1* promoter, thus determining the number of axillary branches. In addition, *APOLO* is involved in leaf hyponasty, likely through the epigenetic regulation of the auxin synthesis-related gene *YUCCA2*, and *PID* and *WAG2*, two genes encoding kinases implicated in the phosphorylation of PIN transporters and auxin redistribution under low R/FR ratio. Furthermore, we developed an approach to assess RNA biological activity based on direct application of RNA to *Arabidopsis thaliana* leaves. Notably, leaves sprayed with *in vitro*-transcribed *APOLO* triggers auxin-related disorders and results in an altered response of plants to the environment. Altogether, our data supports a role for the lncRNA *APOLO* in the epigenetic regulation of SAS and reveals the potential of exogenous lncRNAs as active biomolecules for the modulation of plant-environment interactions.

## RESULTS

### The lncRNA *APOLO* regulates axillary branching

In a previous study, we generated *Arabidopsis thaliana* plants with elevated or reduced levels of *APOLO* (Ariel et al., 2014, Ariel et al 2020, Fonouni-Farde et al., 2022). When growing these plants, we observed an abnormal number of axillary branches. To investigate this observation, we grew side-by-side wild-type (WT) plants and plants from two independent *APOLO* overexpressing lines (*35S:APOLO1* and 2) and two knockdown lines (*RNAi APOLO1* and 2,) and two CRISPR/Cas9-mediated deletion lines (*CRISPR APOLO* 1 and 2) and quantified the number of branches 15 days after bolting. Interestingly, all tested lines exhibited a higher number of axillary branches than WT plants (Figure 1A). By analyzing the plants bearing the entire intergenic region between *PID* and *APOLO* directing the expression of the reporter gene *GUS*, we determined that this promoter region is inactive in axillary buds, in contrast to the leaf petiole where the activity of the *APOLO* promoter is observed (Figure 1B). Accordingly, *APOLO* transcript levels were higher in leaves than in samples enriched in axillary buds, and in both samples expression levels are higher than in roots (Figure 1C).

**Figure 1.**
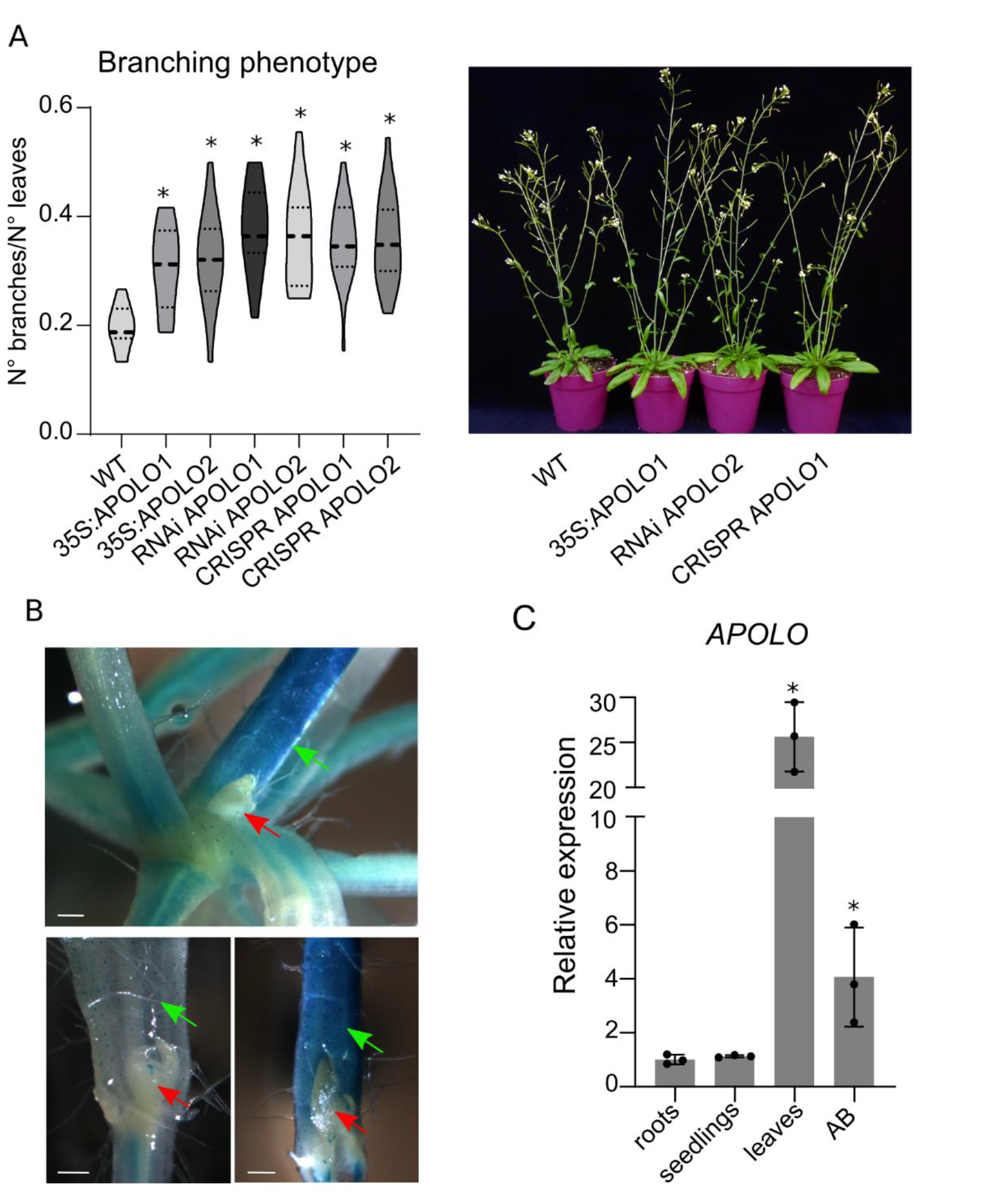
*APOLO* deregulation affects branching. **A.** Number of branches/leaves ratio, 15 days after bolting using 2 independent lines of 35S:*APOLO* (OE), 2 of RNAi and 2 CRISPR plants. Asterisks (*) indicate Student’s t test P≤ 0.05 (n=24) between each line and WT. Photograph of the plants. **B.** GUS staining of *proAPOLO:GFP–GUS* plants. Red arrows indicate axillary buds, while green arrows indicate petioles. White lines correspond to 100μm. **C.** Relative *APOLO* transcripts levels measured by RT-qPCR in different organs of WT plants in control conditions. AB means axillary buds-enriched sample. All the individual measures are shown with black dots. Asterisks (*) indicates Student’s t test P≤ 0.05 (n=3) between each organ and roots, which was the first organ where *APOLO* was measured (Ariel et al., 2014).

### The lncRNA *APOLO* directly controls the expression of the branching master regulator *BRC1*

Previous studies have shown that *APOLO* recognizes a plethora of auxin-responsive genes by sequence complementarity, thus forming R-loops (Ariel et al., 2020). Thus, we searched for genes among *APOLO* targets (Ariel et al., 2020) that could explain the observed differences in branching. Among them, we identified *BRC1*, a known regulator of branching in *Arabidopsis thaliana* (Aguilar-Martínez et al., 2007). To assess if *APOLO* plays a role in modulating *BRC1* we performed RT-qPCR using *APOLO* deregulated plants. This analysis revealed that *BRC1* basal transcript levels were much lower in axillary buds of plants that ectopically express or lack *APOLO* expression (Figure 2A). We then interrogated publicly available epigenomic datasets (Veluchamy et al., 2016; Xu et al., 2017; Ariel et al., 2020) and found that the epigenetic profile of the *BRC1* locus resembles a typical *APOLO* target: high H3K27me3 deposition, LHP1 binding, *APOLO* interaction and a R-loop formation (Figure 2B). Notably, we found that *APOLO* binds 6,277 bp upstream of *BRC1* near the 5’ end of the neighboring gene. We validated the interaction between *APOLO* and this genomic element using ChIRP-qPCR and DRIP-qPCR in WT vs. *CRISPR APOLO* 1 seedlings, we confirmed that *APOLO* interacts with the *BRC1* locus and demonstrated that the coincident R-loop is mediated by *APOLO* (Figure 2C and D). Since the epigenomic data was generated in developing seedlings, we performed ChIP-qPCR to determine the H3K27me3 profile of the *BRC1* locus in mature leaves and axillary buds. Our data shows that the levels of H3K27me3 are higher in leaves than in buds (Figure 2E); in agreement with the *BRC1* expression in axillary meristems (Aguilar-Martinez et al., 2007). Notably, *BRC1* repression in leaves and axillary buds contrasted with the expression of *APOLO* in these tissues (Figure 1C). Remarkably, *BRC1* expression was partial repressed in *APOLO* deregulated lines (Figure 2A) and correlated with an enhanced deposition of H3K27me3 in axillary buds (Figure 2F). Taken together, our data suggest that *APOLO* participates in the epigenetic regulation of *BRC1*.

**Figure 2.**
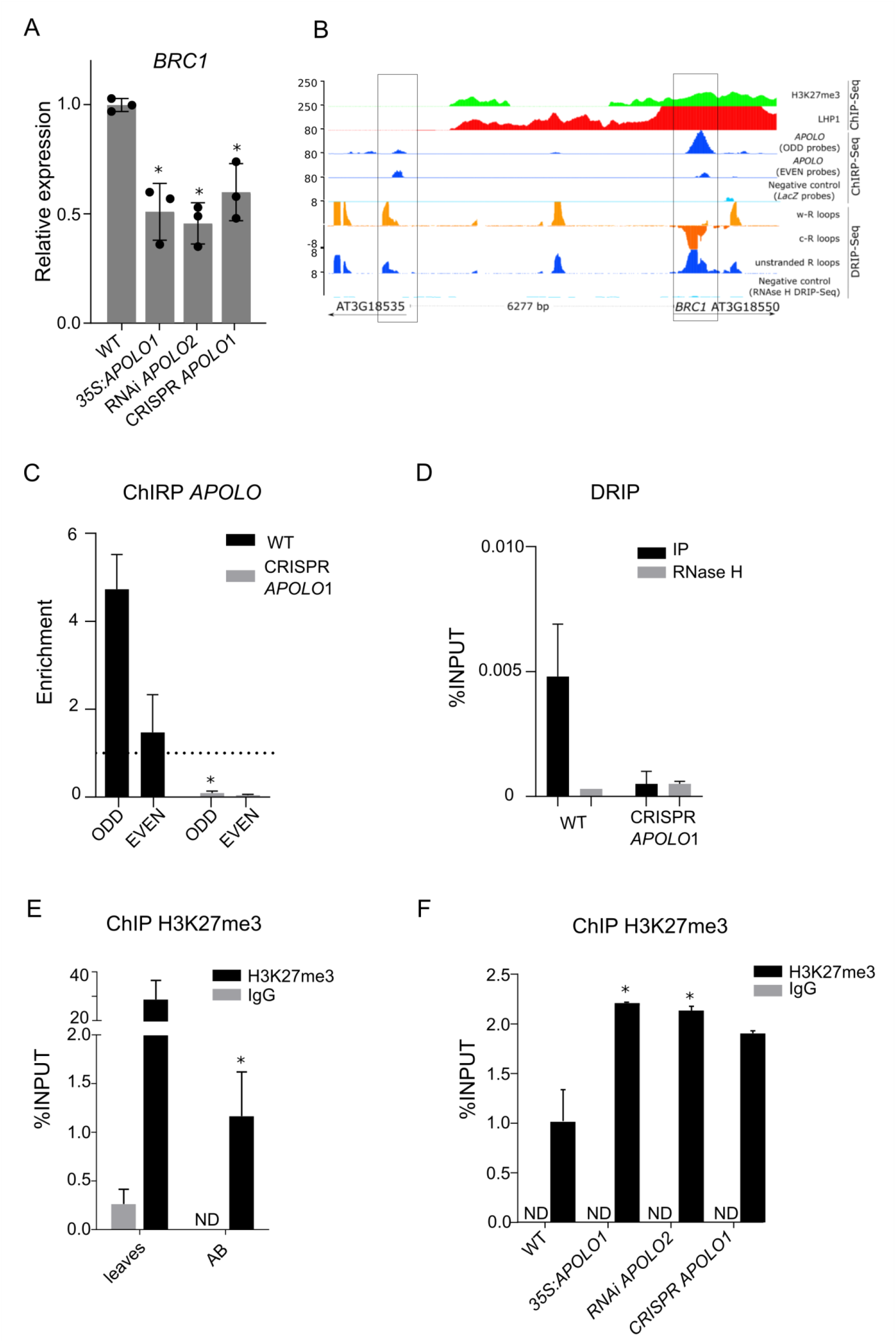
Epigenetic regulation of *BRC1* is mediated by *APOLO*. **A.** Relative *BRC1* transcripts levels measured by RT-qPCR in axillary bud-enriched samples of *APOLO* deregulated lines. All the individual measures are shown with black dots. Asterisks (*) indicate Student’s t test P≤0.05 (n=3) between *APOLO* lines and WT. **B.** Epigenomic landscape of *BRC1* locus. Track 1: H3K27me3 deposition by ChIP-seq. Track 2: LHP1 deposition by ChIP-seq. Tracks 3-5: *APOLO* recognition by ChIRP-seq (tracks 3 and 4, using ODD and EVEN set of probes against *APOLO*, respectively; track 5 negative control using LacZ probes). Tracks 6-9: R-loops formation by DRIP-seq on Watson strand (track 6), Crick strand (track 7), or unstranded sequencing (track 8). DRIP negative control after RNAseH treatment is shown in track 9. Gene annotation is shown at the bottom. **C.** *APOLO* interaction by ChIRP-qPCR in WT vs. CRISPR *APOLO*1. Asterisk indicates Student’s t test P≤0.05. The mean of ODD and EVEN probes ChIRP-qPCR is expressed (value 1 is background level, defined by LacZ probes ChIRP). **D.** R-loop formation by DRIP-qPCR in WT vs. CRISPR *APOLO1*. **E.** H3K27me3 deposition over the *BRC1* locus determined by ChIP-qPCR in leaves vs. axillary bud-enriched samples of WT plants. Asterisk indicates Student’s t test P≤0.05 between the IP and the control with IgG. **F.** H3K27me2 deposition over the *BRC1* locus determined by ChIP-qPCR in axillary bud-enriched samples of 35S *APOLO*1, RNAi *APOLO2* and CRISPR *APOLO1* plants. Asterisks indicate Student’s t test P≤0.05 for each line between the IP and the control with IgG.

### A tissue-specific chromatin loop encompassing *BRC1* promoter depends on *APOLO* transcript levels

It has been reported that *APOLO* modulates chromatin 3D conformation upon recognition of target genes (Ariel et al., 2020). Thus, we profiled chromatin loops identified by capture-HiC in shoots and roots (Huang et al., 2021). This analysis revealed a chromatin loop linking *BRC1* and its neighboring gene, precisely at the *APOLO* binding sites we identified (Figure 3A). This chromatin loop is found primarily in leaves and is dependent on CLF activity, suggesting that H3K27me3 acts as a key regulatory feature for chromatin dynamics at this locus. We then wondered how local chromatin conformation at this locus was organized in axillary buds. To address this caveat we performed 3C-PCR and amplicon sequencing. Remarkably, this analysis revealed that the chromatin loop at the *BRC1* locus is only formed in leaves (Figure 3B) and correlates with high *APOLO* expression and *BRC1* repression. Then, we performed 3C-PCR amplicon sequencing of axillary buds from *APOLO* deregulated plants. Notably, this analysis revealed that the interaction identified in leaves of WT plants was also found in axillary buds of plants over expressing *APOLO* (Figure 3C). However, in *APOLO* knockdown or deletion plants this interaction was not found in axillary buds (Figure 3C). Collectively, these results indicate that *APOLO* transcripts are necessary and sufficient to modulate the chromatin environment of *BRC1* and fine-tune its expression in different tissues.

**Figure 3.**
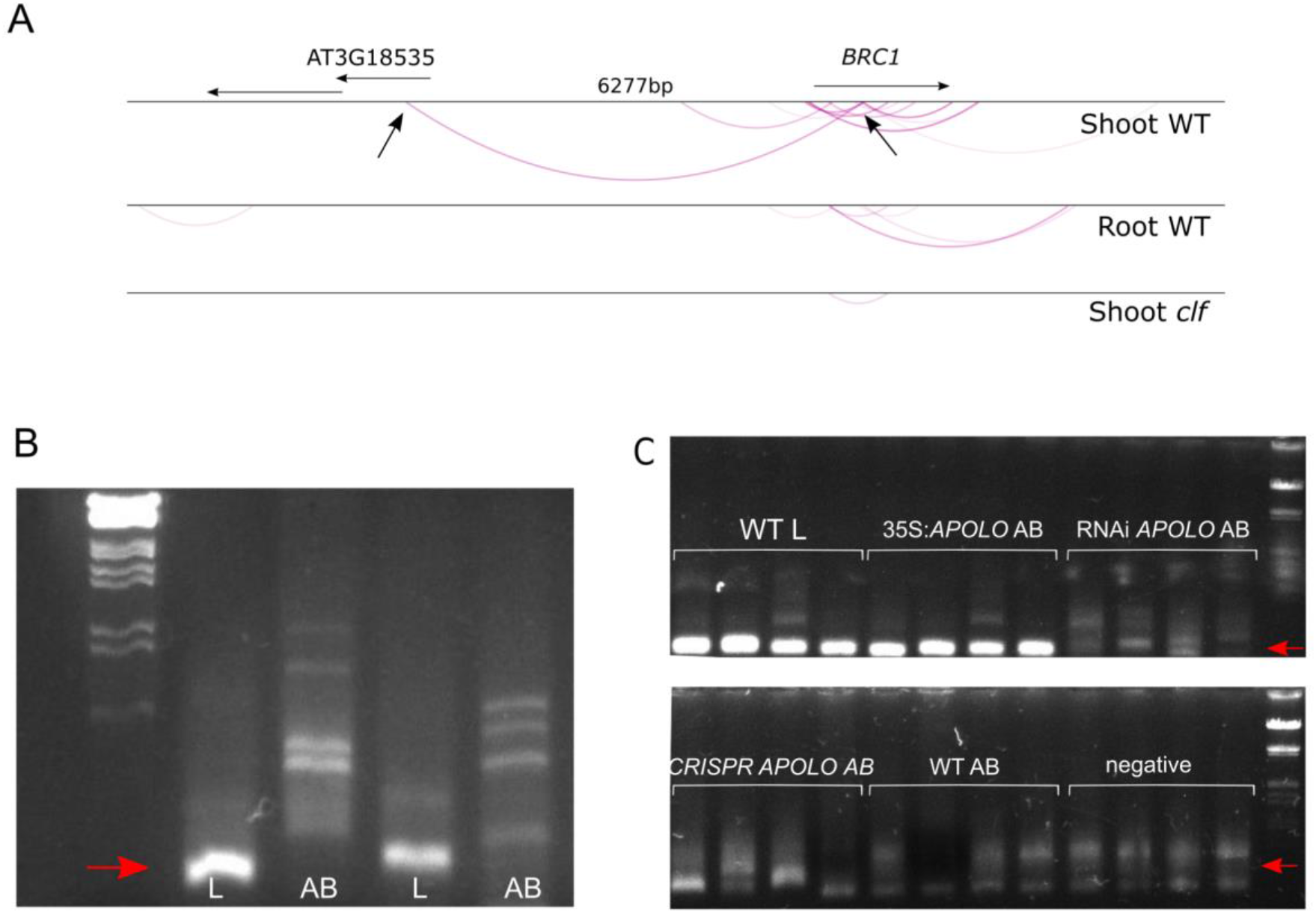
A shoot-specific PRC2-dependent chromatin loop encompasses the intergenic region between *BRC1* and its upstream neighbor *AT3G18353*. **A.** Capture HiC in shoots and roots of WT and shoots of *clf* plants. Track 1 shows gene annotation. Track 2, chromatin loops in *BRC1* locus of shoot WT plants identified by Capture-HiC. Black arrows indicate the sites of *APOLO* binding shown in Figure 2. Track 3, chromatin loops in the *BRC1* locus of roots WT. Track 4, chromatin loops in the *BRC1* locus of *clf* shoots. **B.** 1% agarose gel photograph. 3C-PCR in leaves (L) vs. axillary bud-enriched sample (AB) of WT plants. Two replicates are shown. Red arrow indicates the bands corresponding to the chromatin loop relegation and amplification. **C.** 1% agarose gel photograph. 3C-PCR in WT leaves (WT L) vs axillary bud-enriched samples of WT (WT AB), *35S:APOLO1 (35S:APOLO* AB), RNAi *APOLO2* (RNAi *APOLO* AB), CRISPR *APOLO1* (CRISPR AB) and WT (WT AB). Four replicates are shown. Arrows indicate the bands corresponding to the chromatin loop religation and amplification.

### A low R/FR light ratio modulates chromatin loop formation in the *BRC1* locus

In order to decipher what developmental or environmental cues that influence local *BRC1* chromatin dynamics, we first screened which pathways know to affect *BRC1* expression regulate *APOLO* transcript levels in axillary buds. To this end, we performed exogenous treatments with the phytohormones auxin and strigolactone, exposing plants to a low R/FR light ratio and in plants with altered sugar metabolism (Schluepmann et al., 2003; Figure 4A). In agreement with previous reports *APOLO* was induced by auxin (Ariel et al., 2014 and 2020). Interestingly, its transcript levels in axillary buds decreased in response to 2 h of low R/FR ratio. In contrast, no regulation was observed by strigolactone or in plants with altered sugar metabolism. Considering the dynamic behavior of *APOLO* in response to auxin (Ariel et al., 2014), we characterized its response under low R/FR growth conditions. *APOLO* transcript levels in axillary buds first decreased and gradually increased to basal levels, however, *BRC1* transcripts exhibited the opposite behavior (Figure 4B). We then characterized axillary branching in response to a low R/FR treatment after bolting. Plants over expressing *APOLO* had a higher number of branches than WT both, in WL and also in response to a low R/FR, and both genotypes showed a reduced number of branches in low R/FR. However, we observed a trend where *APOLO*-RNAi and CRISPR plants did not display a full response to low R/FR, and produced a similar number of branches in both light treatments (Figure 4C). Considering the key role of H3K27me3 in chromatin loop formation (Figure 3A), we wondered whether this mark was modulated by low R/FR ratio. We performed ChiP-qPCR on axillary buds enriched samples and found normal H3K27me3 levels in control vs 2 h of low R/FR (Figure 4D). However, after 8 h of low R/FR treatment, when *APOLO* levels returned to basal levels, the chromatin loop observed in leaves was also detectable in axillary buds (Figure 4E). This result agrees with our previous observations that *APOLO* over expression induces chromatin loop formation (Figure 3C). Altogether, our results indicate that *APOLO* can rapidly fine-tune chromatin loop dynamics over the *BRC1* locus although H3K27me3 levels remain constant, hinting at a lncRNA-mediated mechanism alternatively controlling gene expression in different cell types. Low R/FR ratio dynamically represses *APOLO* expression and induces *BRC1*. Considering the significant role of *APOLO* in *BRC1* chromatin loop formation, our results points to a role of this noncoding transcript in the epigenetic regulation of *BRC1* in response to low R/FR ratio.

**Figure 4.**
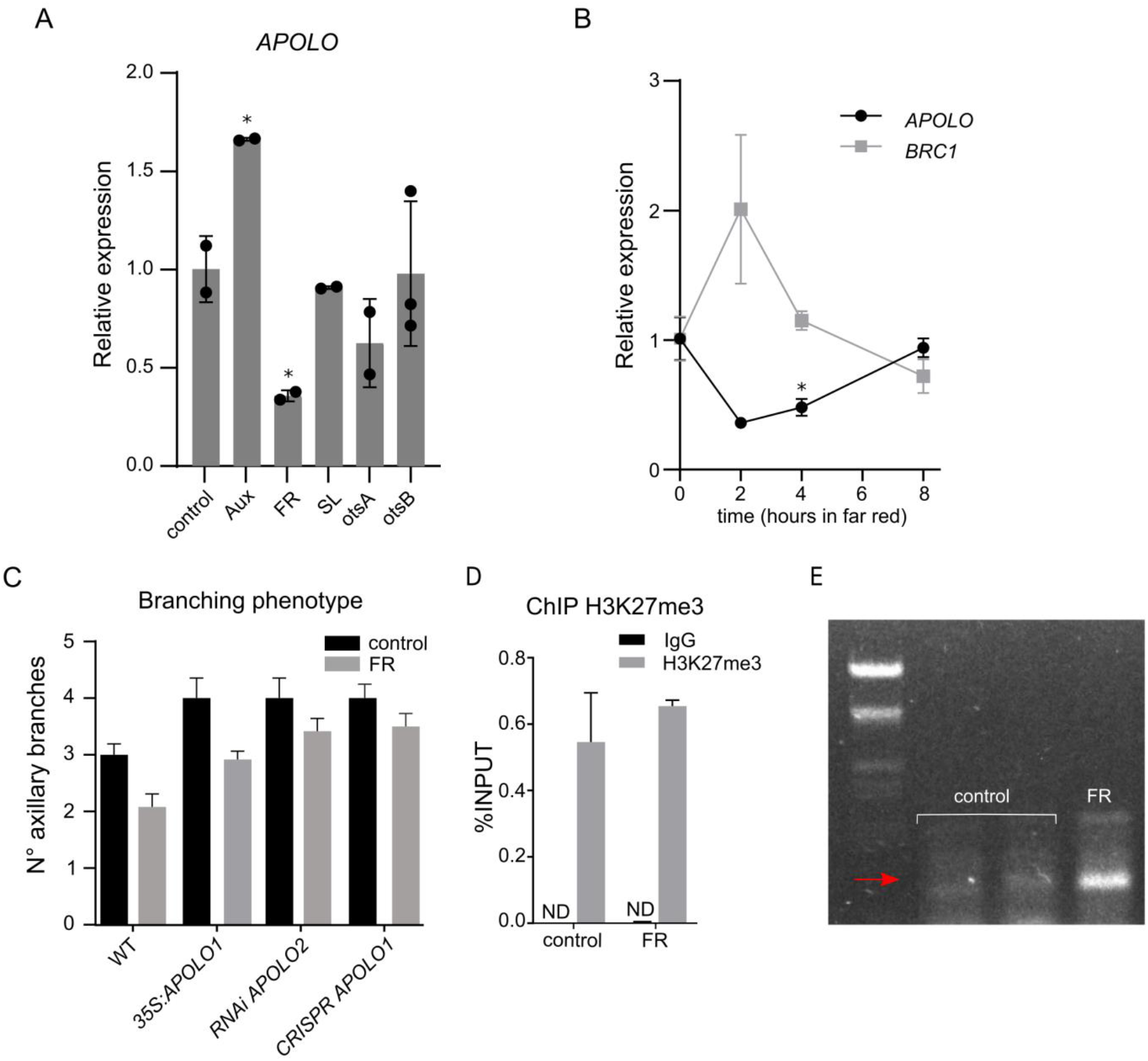
*APOLO* regulates *BRC1* expression under far-red light conditions. **A.** Relative *APOLO* transcripts levels measured in axillary bud-enriched samples by RT-qPCR in WT plants without treatment (control), treated with auxin (Aux), exposed to WL+FR (FR), with strigolactone application (SL) and in axillary bud-enriched samples of otsA (OE T6P synthase, plants that accumulate more trehalose 6 phosphate) and otsB plants (OE T6P phosphatase, accumulating less trehalose 6 phosphate). Asterisks indicate Student’s t test P≤0.05 (n=3) between the treatments and control. **B.** Relative *APOLO* and *BRC1* transcripts levels measured by RT-qPCR in axillary bud-enriched samples of WT plants at 0, 2, 4 and 8h of FR light treatment. **C.** Branching phenotyping. Number of branches after 15 days in control conditions and with WL+FR (FR). **D.** H3K27me3 deposition over the *BRC1* locus determined by ChIP-qPCR in axillary bud-enriched samples after 8h of white light supplemented with far-red light vs. white light. ND means non detected. **E.** 1% agarose gel photograph. 3C-PCR in WT axillary bud-enriched samples with and without far-red light supplementation. The arrow indicates the band corresponding to the chromatin loop religation and amplification.

### *APOLO* participates in the hyponastic response of leaves to low R/FR by modulating auxin homeostasis

Having established that a low R/FR light ratio modulates *APOLO* expression, we investigated whether *APOLO* is involved in the control of leaf hyponasty, which is another hallmark of SAS in *Arabidopsis thaliana*.

Previous studies have shown that *APOLO* directly regulates in *cis* its neighboring gene *PID* (Ariel et al., 2014) and in *trans* the *PID* homolog *WAG2* (Ariel et al., 2020). *PID* and *WAG2* encode two kinases in charge of determining the position of PIN auxin transporters in the cell membrane, thus modulating auxin efflux (Benjamins et al., 2001; Dhonukshe et al., 2010). In addition, other studies have shown that *APOLO* also controls the expression of *YUCCA2* by coordinating the action of LHP1 and VIM1 on histone and DNA methylation (Fonouni-Farde et al., 2022). Among other *YUCCA* genes, *YUCCA2* is a key factor in auxin synthesis (Mashiguchi et al., 2011). Considering that auxin synthesis, accumulation and distribution (Keuskamp et al., 2010; de Wit et al., 2015; Michaud et al., 2017; Pantazopoulou et al., 2017; Küper et al., 2023) control hyponasty in response to a low R/FR ratio, we hypothesized that *APOLO* is involved in this response. To test this idea, we exposed plants to white light supplemented with FR (low R/FR ratio) and took pictures of plants every 30 min during an 8 h-treatment period, which allowed us to measure the angle of the leaf exhibiting the highest upward movement by the end of the experiment. The same leaf from each plant was considered throughout the entire experiment. In WT plants, the maximal angle was reached after 4 h of treatment, ranging from 30 to 40 degrees. In contrast, *APOLO* knockdown and knockout plants exhibited a delayed upward movement and had a final leaf angle ranging from 20 to 30 degrees. Finally, plants that overexpress *APOLO* displayed the most drastic effect in terms of delay and the final leaf angle, which did not exceed 20 degrees (Figure 5A). Collectivelly, our results indicate that in addition to branching, deregulation of *APOLO* compromises the hyponastic response to low R/FR ratio.

**Figure 5.**
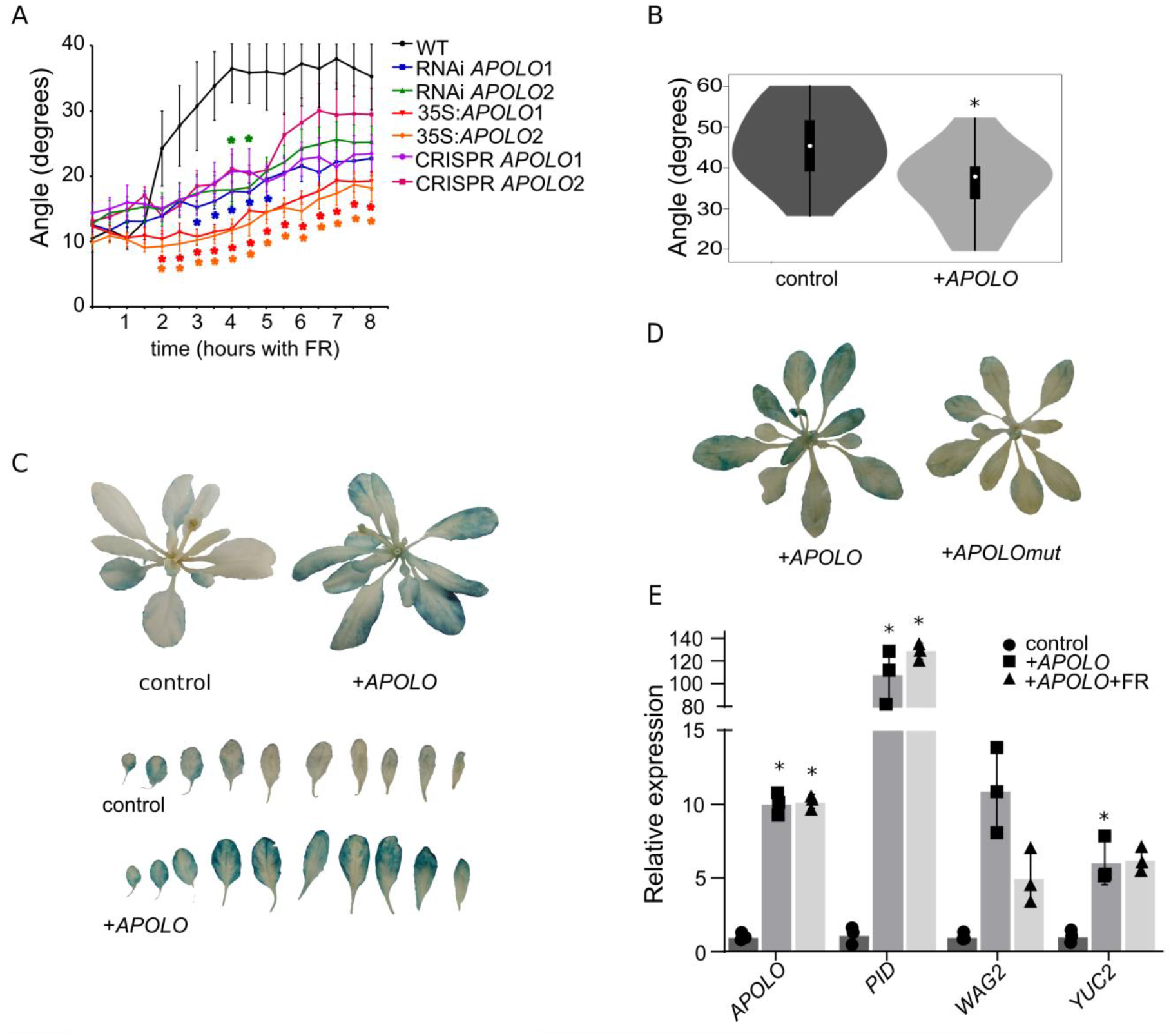
*APOLO* is involved in low R/FR dependent hyponasty. **A.** Angle of elevation of rosette leaves when plants are exposed to WL+FR during 8 h. **B.** Angle of elevation of rosette leaves when WT plants are sprayed with *APOLO* RNA vs. *GFP* RNA transcribed *in vitro* after 8 h exposure to WL+FR. **C.** GUS staining of *DR5:GUS* plants sprayed with *GFP* and *APOLO* RNAs transcribed *in vitro* after 8 h exposure to WL+FR. Individual leaves are shown in the lower panel. **D.** GUS staining of *DR5:GUS* plants sprayed with *APOLO* and *APOLO*mut (i.e. with abolished R-loop formation) RNAs transcribed *in vitro* after 8 h exposure to WL+FR. **E.** Relative *APOLO, PID, WAG2* and *YUC2* transcripts levels in nucleus of WT plants without treatment (control), WT sprayed with *APOLO* RNA transcribed *in vitro* after 8 h exposure to WL (+*APOLO*) and WL+FR (+*APOLO*+FR). Asterisks indicate Student’s t test P≤0.05 (n=3) for the different genes between each treatment and control.

### Exogenous application of *in vitro*-transcribed *APOLO* alters the plant auxin homeostasis and the response to low R/FR ratio

The effect of *APOLO* over expression led us to wonder if an exogenous treatment with *in vitro*-transcribed *APOLO* could modulate the response of the plant to the environment. It has been shown that plants can absorb exogenous RNAs, which have been increasingly used as double-stranded transcripts triggering the production of small RNAs capable of silencing endogenous genes or blocking the infection of viruses, fungi or even insects (Rodriguez et al., 2023). However, the potential use of epigenetically active lncRNAs as exogenous bioactive molecules remains to be demonstrated. To test this hypothesis, we transcribed *APOLO in vitro* and sprayed *Arabidopsis thaliana* plants one day before treatment with WL+FR. We found that after 8 h treatment, the final angle of the highest leaf was significantly lower in plants sprayed with *APOLO* RNA than in plants sprayed with *GFP* RNA used as a mock control (Figure 5B). To determine if the plant differential behavior under *APOLO* exogenous treatment was related to auxin homeostasis, we also sprayed *DR5:GUS* plants that one day later were subjected to low R/FR light treatment. Strikingly, only the plants sprayed with *APOLO* RNA exhibited expanded staining at the contour of the leaf blade, indicating a drastic impact on auxin synthesis, distribution and/or signaling (Figure 5C). To test if this effect is mediated by direct interaction of *APOLO* RNA to chromatin, we used a mutagenized *APOLO RNA* (*APOLOmut*) lacking two TTCTTC boxes that are known to be essential for R-loop formation and target recognition (Ariel et al., 2020). Remarkably, *APOLOmut RNA* was unable to induce the expression of the auxin reporter, indicating that lncRNA-DNA interaction was required to trigger a biological response. To validate these results, we extracted nuclei from leaves sprayed with either *APOLO* or *GFP* to quantify transcript levels. This analysis revealed an increase in levels of full-length *APOLO* in the nuclei by sprayed with *in vitro*-transcribed *APOLO* (Figure 5E). Considering the link between SAS and auxin homeostasis, and the control of *APOLO* over auxin related genes, we wondered how these key genes responded in sprayed plants. Interestingly, the abundance of *PID, WAG2* and *YUCCA2* transcripts was enhanced, indicating that higher levels of *APOLO* triggered the transcription of target genes and ultimately deregulating auxin homeostasis. Altogether, our data shows that exogenously applied lncRNAs are sufficient to trigger a specific epigenetic-mediated activation of genes modulating auxin homeostasis and coordinating SAS.

## DISCUSSION

Light is indispensable for plant growth and developmental plasticity allowing these sessile organisms to optimize light capture and utilization. Physiological and molecular analyses in *Arabidopsis thaliana* and other plant species indicate that plastic responses to changes in light intensity and quality relay on the integration of signals from a broad range of hormonal players (Fernandez-Milmanda and Ballaré, 2021) with auxin signaling playing a key role in SAS responses (Tao et al., 2008; Li et al., 2012). In this context, the plant nucleus provides a major hub for signal integration at the chromatin level (Patitaki el at., 2022), including the re-organization of the genetic information in three dimensions. Here, we show that the lncRNA *APOLO* participates in SAS by modulating local chromatin conformation of distinct subsets of target genes in axillary buds and leaves, modulating auxin homeostasis and coordinating SAS. *APOLO* differential transcriptional levels in these organs fine-tune the expression of key genes involved in SAS responses. It was previously reported that *APOLO* recognizes multiple auxin-responsive loci *in trans* across the *Arabidopsis thaliana* genome through sequence complementarity, forming R-loops. As a result, *APOLO* impacts target chromatin loop dynamics and affects gene transcriptional activity (Ariel et al 2020). More recently, the link between lncRNA-mediated R-loop formation and chromatin 3D conformation dynamics was also uncovered in mammalian cells (Luo et al., 2022), pointing to conserved mechanisms both in plants and animals, linking noncoding transcription, non-B DNA structures, and gene activity.

Our work reveals that *APOLO* is expressed at low levels in axillary buds, in contrast to its higher expression in leaves. In response to low R/FR, the expression of *APOLO* and *BCR1* follow an opposite pattern suggesting that FR may enhance *BCR1* expression by alleviating the repression caused by the *APOLO*-dependent chromatin loop in the *BCR1* locus. Interestingly, in the *APOLO* RNAi and CRISPR lines, FR supplementation appeared to have an attenuated effect in the suppression of branching, suggesting that, in the long term, the *BCR1* mis-regulation imposed by the loss of *APOLO* could alter the plastic responses to shade.

Among angiosperms, the suppression of bud outgrowth depends on a highly conserved mechanism that involves *BRC1*-like genes. In *Nicotiana tabacum*, a subset of lncRNAs have been proposed to be involved in axillary bud outgrowth (Wang et al., 2022). At least four lncRNAs were proposed to act downstream *NtTB1* (the *Nicotiana* ortholog of *BRC1*), whereas MSTRG.28151.1 was identified as an antisense lncRNA of *NtTB1*. MSTRG.28151.1 knockdown significantly attenuated *NtTB1* expression and resulted in larger axillary buds. However, the molecular mechanism involving the natural antisense transcript of *NtTB1* remains uncertain. In maize leaves, several chromatin loops were identified over the *TB1* locus in agreement with H3K27me3 deposition, hinting at an epigenetic silencing of *TB1* in these organs (Ricci et al. 2019). Nevertheless, it remains unknown whether H3K27me3 deposition and/or 3D chromatin conformation dynamics of the *TB1* locus are controlled by lncRNAs. Here we show how an intergenic lncRNA regulates axillary branching in *Arabidopsis thaliana* by directly recognizing the *TB1* ortholog, *BRC1*, in *trans. APOLO* deregulation affected H3K27me3 deposition, although higher levels of *APOLO* induced by low R/FR seem to boost the formation of a chromatin loop encompassing *BRC1* promoter and transcriptional repression, independently from H3K27me3 deposition.

In addition, our observations implicate *APOLO* in the hyponastic response to a low R/FR ratio through the modulation of previously reported target genes. It is known that FR modulates auxin synthesis (Michaud et al., 2017), dynamics (Pantazopoulou et al., 2017) and redistribution (Keuskamp et al., 2010; Küpers et al., 2023), which are necessary to trigger leaf hyponasty. FR inactivates phyB, which results in increased activity of the key transcription factor PHYTOCHROME INTERACTING FACTOR7 (PIF7) (Li et al., 2012). PIF7 controls PIN and auxin redistribution (Michaud et al., 2017) and, along with other members of the PIF family, regulates the hyponastic response to FR radiation (Küpers et al., 2023). Considering that PIF7 activates auxin synthesis through *YUCCA* genes (Li et al., 2012), and that *APOLO* is induced by auxin, it is possible that PIN redistribution may rely on *APOLO*-mediated epigenetic activation of *PID* and *WAG2* (Ariel et al., 2020), which can phosphorylate PIN proteins in response to FR, triggering hyponasty. Furthermore, *APOLO* directly regulates the auxin synthesis-related gene *YUCCA2* (Fonouni-Farde et al., 2022), pointing to a positive feedback loop mediated by the lncRNA in response to low R/FR ratios. It is known that *APOLO* coordinates histone and DNA methylation to block *YUCCA2* transcription in basal conditions, through direct interaction with LHP1 and the hemi-methylated DNA binding protein VIM1. In response to warmth, *APOLO* levels decrease, the ribonucleoprotein complex is disrupted and *YUCCA2* transcription increases, triggering auxin-dependent thermomorphonesis (Fonouni-Farde et al., 2022). Warm temperatures promote hypocotyl and petiole elongation and leaf hyponasty (Quint et al., 2016; Casal & Balasubramanian, 2019). These growth patterns are similar to those activated during shade avoidance and, in fact, the responses to warm temperatures and shade share some important molecular players. It is known that both stimuli can be perceived by phyB in *Arabidopsis thaliana*, and mediated by PIF transcription factors which will ultimately activate growth-promoting target genes. PIF4 and PIF7 have been shown to be master regulators of thermomorphogenesis (Koini et al. 2009; Quint et al., 2016) and shade avoidance responses (Li et al. 2012), respectively. Notably, recent studies also revealed a major role for PIF7 during thermomorphogenesis (Fiorucci et al., 2020; Chung et al 2020; Burko et al., 2022).

Low R/FR or warm ambient temperature treatments activate the auxin pathway (Kohnen et al., 2016; Ballstaedt et al., 2019) through the PIF-dependent transcriptional regulation of *YUCCA* genes. It was demonstrated that PIF4 modulates hyponastic leaf movement under warm temperature conditions by a two-branched auxin signaling pathway (Park et al., 2019). Thermo-activated PIF4 directly induces *PID* transcription in petiole cells, resulting in polar accumulation of auxin. In another route, the PIF4-YUC8 branch promotes auxin production in the blade, which is transported to the petiole and functions as the substrate for PIN3 machinery. PIF4-mediated *PID* transcription occurs mostly in the abaxial petiole region leading to PIF4-mediated leaf hyponasty (Park et al., 2019). Interestingly, PIF4 recognizes *PID* promoter, i.e. the intergenic region between *PID* and *APOLO. APOLO* deregulation also affects hypocotyl elongation in response to warmth, and the transcriptome of seedlings over-expressing APOLO resemble the one of WT plants grown in warmth (Fonouni-Farde et al., 2022). Our data show that *APOLO* deregulation impacts axillary branching and hyponasty, further supporting the role of *APOLO* in integrating of the auxin-related cross-talk between thermomorphogenesis and SAS.

Recently, FR-regulated lncRNAs were identified in *Dendrobium officinale*, and it was suggested that they might participate in SAS through hormone signal transduction and DNA methylation (Li et al., 2021). However, their potential roles in this pathway remain unknown. In *Arabidopsis thaliana*, the lncRNA *HIDDEN TREASURE 1* (*HID1*) positively regulates photomorphogenesis under low R/FR ratio by downregulating *PIF3* expression levels (Wang et al., 2014). It has been suggested that *HID1* takes part in a large nuclear ribonucleoprotein complex and associates with the first intron of the *PIF3* locus. Another *Arabidopsis thaliana* lncRNA involved in photomorphogenesis is *BLUE LIGHT INDUCED LNCRNA 1* (*BLIL1*), which participates in the regulation of photomorphogenesis under blue light conditions and in response to mannitol stress through miR167 sequestering as a target mimicry lncRNA (Sun et al., 2020). Similarly, our data show that the auxin-responsive lncRNA *APOLO* is also involved in SAS. Interestingly, we uncovered an organ-specific role for *APOLO*, depending on its relative transcriptional accumulation. Furthermore, the exogenous application of *in vitro* transcribed *APOLO RNA* triggers auxin-mediated responses. There is an increasing number of reports in plants indicating that the application of exogenous dsRNAs trigger changes in gene expression, sometimes targeting endogenous or pathogen-specific genes resulting in resistance to infection (Rodriguez Melo et al., 2023). Our data shows that exogenously applied *APOLO* RNA activate the expression of specific targets through direct interaction with chromatin and that this response is underpinned by specific sequence motifs. Our work opens the door to new strategies to modulate plant growth and their response to environmental cues by the exogenous application of epigenetically active lncRNAs.

## METHODS

### Plant material

All lines used are in Columbia-0 background. The *APOLO* overexpressing lines were reported in Ariel et al., 2020 and the RNAi lines in Ariel et al., 2014. The CRISPR *APOLO* 1 line was reported in Fonouni-Farde et al., 2022 and the CRISPR *APOLO* 2 was generated by creating a heritable 450bp deletion using the methodology described by Durr et al., 2018. Probes used are indicated in Supplementary Table 1. The pro*APOLO*:GFP-GUS line was reported in Ariel et al., 2020. otsA and otsB lines were reported in Schluepmann et al., 2003. Plants were grown under long day conditions (16 h light, 90 μmol m-2 s-1 / 8 h dark) at 23 °C.

For SL treatments, 50 μL of 10 μM GR24^5DS^ and 0.1% (v/v) Tween-20 were added to rosette axillary buds, at bolting stage (1cm-1,5cm) and samples were collected 6 h after beginning of the treatment. For Auxin treatments plants of the same developmental stage were decapitated and the cut stump was inserted into an inverted 0.5 mL tube with 500 μL of 0.6% (w/v) agarose and 50 μM NAA or 0.005% NaOH 1N (control), samples were collected 24 h after decapitation. For FR treatments plants of the same developmental stage were exposed to WL+FR (R/FR ratio=0.11; control R/FR=2.7).

### Branching phenotype characterization

Rosette branches (> 0.5 cm) were counted 15 days after the bolting stage. 24 plants of each genotype were analyzed.

For the branching phenotypes in low R/FR, plants were grown in normal conditions (R/FR=2.7) until the inflorescence was visible and then in WL+FR (R/FR=0.11) for 15 days before branches were counted. 12 plants of each genotype were analyzed.

### Hyponasty phenotype characterization

3-week-old plants were exposed to WL+FR during 8 h between 10 h to 18 h (photoperiod from 6 h to 22 h). Photos were taken every 30 min and then the angle between the soil and the highest leaf was measured with ImageJ.

### Histochemical staining

3-week-old plants bearing the *APOLO* promoter controlling the expression of the reporter gene *GUS* were used (Ariel et al., 2020). GUS staining was performed as described by Fonouni-Farde et al., 2022.

### RNA Isolation and DNAse treatment

Samples enriched in axillary buds were obtained from the rosette center of 3-week-old plants with a hole puncher. RNA samples were obtained with Trizol. NEB DNAseI was used according to the manufacturer’s instructions.

### Real-Time PCR

A total of 1 ug of RNA was used for oligo(dT) reverse transcription (SuperScriptII, Invitrogen). RT-qPCR were performed using SsoAdvanced Universal SYBR Green Supermix (Bio Rad) on a StepOne Plus apparatus (Applied Biosystems) using standard protocols (40 to 45 cycles, 60°C annealing) and analyzed using the delta delta Ct method using Actin for normalization. Primers used are listed in Supplementary Table 1. The efficiency of all primers was verified by consecutive dilutions of standardized samples.

### Chromatin Immunoprecipitation (ChIP)

ChIP assays were performed on leaves and axillary buds enriched samples of 3 weeks old plants using anti-H3K27me3 (Diagenode pAb-195-050) and anti-IgG (Abcam ab6702), as previously described (Ariel et al., 2020). Primers used are listed in Supplementary Table 1.

### Chromatin Isolation by RNA Purification (ChIRP)

ChIRP was performed in leaves from 3-week-old plants as described previously (Ariel et al., 2020). Primers used are listed in Supplementary Table 1.

### DNA-RNA duplex Immunopurification assay (DRIP)

For DRIP-DNA-qPCR assay, non-crosslinked leaves from 3-week-old plants were used for nuclei purification. The experiment was performed as described by Fonouni-Farde et al., 2022. Primers used are listed in Supplementary Table 1.

### Chromosome Conformation Capture Assay (3C)

3C was performed basically as previously described (Ariel et al., 2014) starting with two grams of leaves or axillary buds enriched samples. Digestions were performed overnight at 37°C with 400U DpnII (NEB). DNA was ligated by incubation in a shaker at 16°C, 100 rpm, for 5 h in 4ml volume using 100U of T4 DNA ligase (NEB). After reverse crosslinking and Proteinase K treatment (Invitrogen), DNA was recovered by Phenol:Chloroform:Isoamyl Acid (25:24:1; Sigma) extraction and ethanol precipitation. Relative interaction frequency was calculated by qPCR. A region free of DpnII was used to normalize the amount of DNA. Primers used are listed in Supplementary Table 1.

### RNA *in vitro* transcription and spray assay

For in vitro transcription of the *APOLO* and *GFP* RNAs, 1 μg of purified DNA of each template including the T7 promoter at the 5’ end was used following the manufacturer instructions (HiScribe T7 High Yield RNA Synthesis kit, NEB). Primers used are listed in Supplementary Table 1. Then 1 μg of the transcribed RNA in water was sprayed to the plants and the following day they were exposed with WL+FR light during 8 h between 10 h to 18 h (photoperiod from 6 h to 22 h).

For the hyponasty phenotype, the plants were photographed for angle measure.

For GUS staining, after 8 h of exposure to WL+FR the staining was performed.

For transcripts measured, samples were taken and frozen with liquid nitrogen. A nuclei isolation was performed and then transcripts were measured by RT–qPCR.

---

**Supplementary Table 1.**
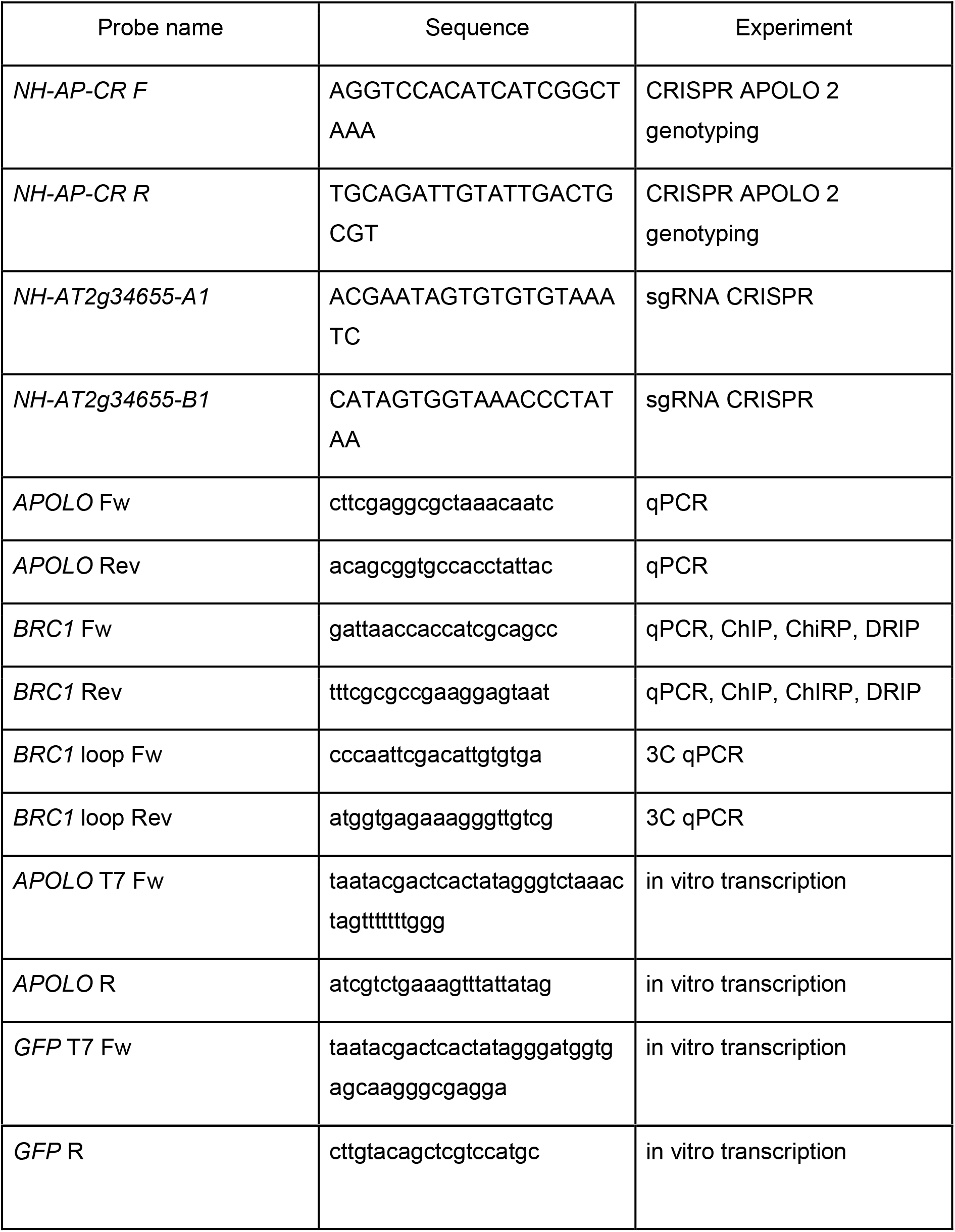
DNA probes used in this project. Their use is indicated in the third column.

## FUNDING

This project was financially supported by grants from Agencia Nacional de Promoción Científica y Tecnológica (PICT), Universidad Nacional del Litoral (CAI+D) and ICGEB CRP-ARG01 (Argentina); International Research Project LOCOSYM (CNRS-CONICET); IPS2 benefits from the support of Saclay Plant Sciences-SPS (ANR-17-EUR-0007); PC has a grant PID2020-112779RB-I00 funded by MCIN/ AEI /10.13039/501100011033 (Spain). LL, PM, CB and FA are members of CONICET. MFM and LF are fellows of the same institution. AM contract PRE2018-083865 was funded by MCIN/AEI/10.13039/501100011033 and FSE invierte en tu futuro.

## DECLARATIONS

### Ethics approval and consent to participate

Not applicable for this study.

### Competing interests

The authors declare that they have no competing interests.

